# Hydraulic constraints of reproduction do not explain sexual dimorphism in the genus *Leucadendron* (Proteaceae)

**DOI:** 10.1101/209460

**Authors:** Adam B. Roddy, Justin van Blerk, Jeremy J. Midgley, Adam G. West

## Abstract

Because of the importance of reproduction in plant life history, the physiological costs of reproduction often influence vegetative structure and function. In dioecious species, these effects can be quite obvious, as different costs of male and female reproductive functions are entirely separated among different individuals in a population. In fire-prone ecosystems, in which recruitment is driven by fire frequency, many plants will maintain their seeds in the canopy, only to be released after a fire. The dioecious genus *Leucadendron* is a notable case of this, as females can maintain their seed cones for years, and, even more interestingly, species in the genus differ substantially in the degree to which males and females are sexually dimorphic. A recent study (Harris and Pannell 2010) argued that the hydraulic costs of maintaining seed cones for many years would effect the degree of sexual dimorphism among species. However, this assumed that shoot hydraulic architecture would be related to traits exhibiting sexual dimorphism. Here we explicitly test this hypothesis on two *Leucadendron* species. We found (1) that metrics of branch ramification used in the previous study to characterize dimorphism do not conform to known scaling relationships and (2) that sexual dimorphism in shoot architecture has no effect on hydraulic efficiency. Both of these results seriously question the pattern described by Harris and Pannell (2010) and suggest that the hydraulic costs of prolonged seed retention in *Leucadendron* do not significantly affect branch architecture.

## Introduction

The extent to which the costs of reproduction in dioecious species are different for male and female plants is a controversial topic in plant evolution (Barrett & Hough 2013). Generally it is thought that costs are higher for females, which has consequences on vegetative structures, creating dimorphism in secondary sexual characters not directly related to reproduction (Dawson & Geber 1999). Dimorphism can be visibly subtle, involving only physiological traits (Dawson & Ehleringer 1993), or more obivous, such as in the case of the genus *Leucadendron* (Proteaceae), native to the Western Cape of South Africa. *Leucadendron* species display a full range of phenotypes, from monomorphism to dimorphism, in leaf size, branch size and ramification, and inflorescence size, with males having smaller, more numerous leaves and branches (Williams 1972; Bond & Midgley 1988; Midgley & Bond 1989; Midgley 2010). Unlike males, females incur the costs of producing nutrient-rich seeds. Furthermore, females of many *Leucadendron* species maintain their cones in the canopy for several years, a strategy termed ‘serotiny’. Serotiny allows seeds to be dispersed only after fire, but preventing the cones from opening during the hot, dry Mediterranean summers requires a continuous supply of water and carbon. Although the water costs of maintaining cones are lower than originally thought (Cramer & Midgley 2009), the relative costs of reproduction are assumed to be higher for females than for males (but see Bond & Maze 1999), and these costs are further expected to increase with increasing serotiny.

The most striking differences between males and females of dimorphic *Leucadendron* species are differences in leaf size and branch ramification (Figure 1). Theoretical and empirical studies in other systems have long considered leaf size to be mechanistically linked to branch size because of the biomechanical constraints on the distribution of leaves (Corner 1949; Ackerly & Donoghue 1998; Olson, Aguirre-Hernández & Rosell 2009). In dimorphic *Leucadendron* species, females have larger leaves and thicker, less ramified stems than their conspecific males. Recently Harris and Pannell (2010) argued that the cause of larger leaves in serotinous females is because of selection on stem hydraulic architecture. Specifically, they argued that thicker branches with fewer nodal junctions are more hydraulically efficient, which is particularly important for serotinous females because maintaining cones requires a continuous supply of water. Implicit in their argument is (1) that water use efficiency should differ among males and females, particularly in highly dimorphic species and (2) that branching reduces hydraulic efficiency. Yet, water use efficiency, as measured by *δ*^13^C isotopes, does not differ between males and females in a diversity of species (Midgley 2010). Further, although branch nodes may have high hydraulic resistance (Zimmermann 1978; Ewers & Zimmermann 1984; Tyree & Alexander 1993; Slingsby 2004), the effects of branching on the entire shoot hydraulic network are unclear.

**Figure 1.**
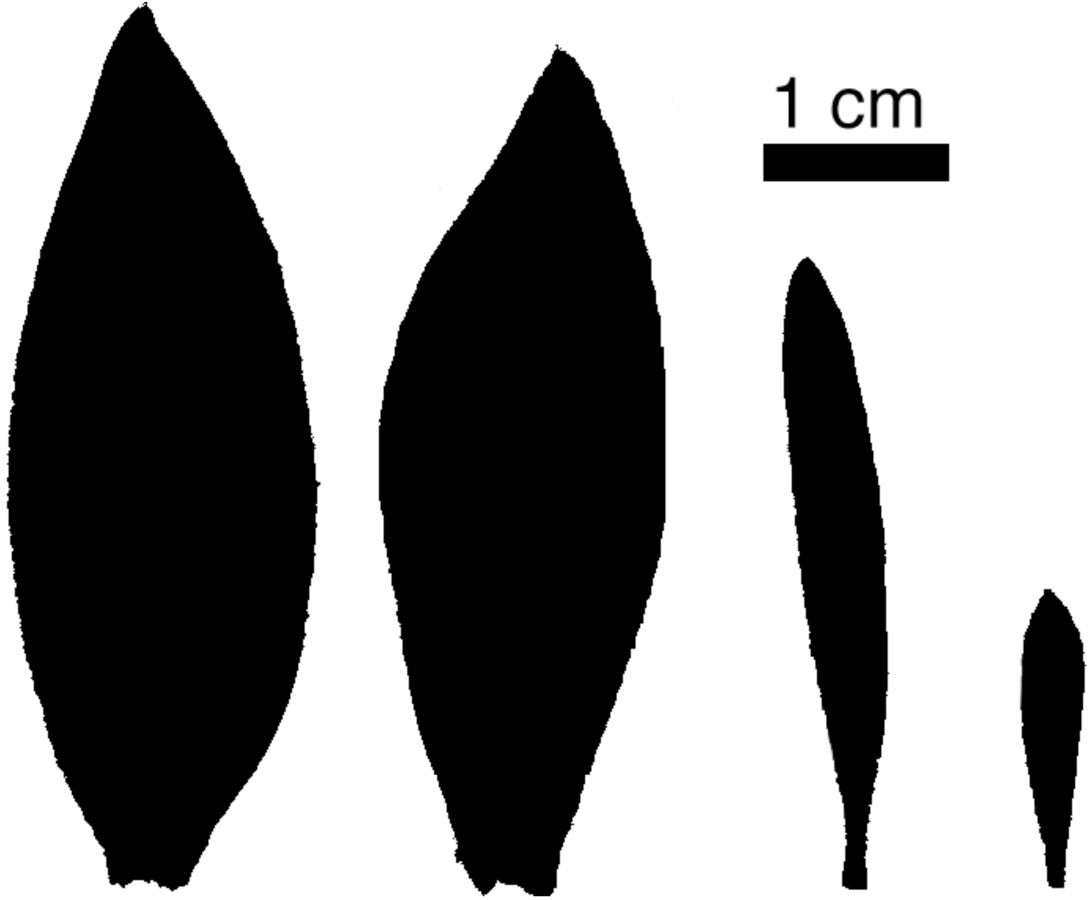
Representative leaves with areas approximating the average leaf area for each group. From left to right: *L. daphnoides* female and male, *L. rubrum* female and male.

To explore possible tradeoffs between shoot architecture and shoot hydraulics, we examined relationships between morphological and physiological traits at three levels: (1) among all individuals (regardless of species and sex), (2) within species (regardless of sex), and, (3) within each species x sex group separately. Some relationships were predicted to be significant at one level but not at other levels. Relationships pertaining to Corner’s scaling rules (e.g. between shoot ramification and leaf area) were predicted to be significant across all species x sex combinations, while other relationships were predicted to be significant within species but with species-specific intercepts (e.g. between ramification and average leaf size). More apropos to this study, correlations with hydraulic efficiency were expected to be significant within species, as the developmental constraints on cell size and cell packing that influence hydraulic efficiency should be common to both males and females (Simonin & Roddy 2017).

Consideration of the assumptions and predictions in the study of Harris and Pannell (2010) depends on formally clarifying a definition of efficiency. In their study, larger plants (e.g. females with thicker stems) were predicted to have higher hydraulic conductance and thus move water more efficiently. These two qualities are not equivalent and not necessarily related. That larger plants move more water would not at all be surprising, as they must support, by definition, a larger body size and leaf area. Hydraulic *efficiency,* however, may differ regardless of plant size. Hydraulic efficiency, in our understanding, refers to the flux of water per investment cost of moving that water. In this case, this would be the hydraulic conductance divided by plant size–while larger plants are expected to move absolutely more water, whether they move more water per unit biomass (or size) investment may differ substantially. For conspecific males and females, there may be differences in hydraulic efficiency, as has been shown in other dioecious species (e.g. Dawson & Ehleringer 1993), but given that there is no strong habitat segregation among conspecific males and females in *Leucadendron,* that the hydraulic costs of bearing cones are relatively small (Cramer & Midgley 2009), and that *δ*^13^C does not differ among males and females, we did not anticipate that differences in hydraulic efficiency between conspecific males and females would be large. Yet, there may nonetheless be relationships between hydraulic architecture and ramification within groups, and we expected these relationships to exist within species with conspecific males and females falling along a single scaling relationship. Despite our prediction that conspecific males and females should exhibit the same scaling relationships, in our analyses we show significant relationships, where they exist, within each species x sex combination in order to be as unbiased as possible in our testing of the predictions of Harris and Pannell (2010).

Regardless of these predictions, how best to characterize and interpret tradeoffs is of significant concern. Tradeoffs between two alternative strategies are thought to exhibit a clear, linear (or near linear) relationship (Figure 2a). In reality, however, few such relationships are this clean. Many bivariate relationships exhibit an upper or lower bound with points occurring in any region below or above this bound (Figure 2b,c, respectively). For example, a hard tradeoff between xylem hydraulic efficiency and safety from embolism has long been thought to be fundamental to hydraulic architecture (i.e. Figure 2a), but the most comprehensive analysis of this relationship reveals that the pattern more resembles that represented in Figure 2b (Gleason *et al.* 2015). Grubb (2016) has argued that while many authors consider such relationships to indicate a tradeoff, this argument is weak. In contrast to a hard tradeoff (Figure 2a), in which any quantile of the data exhibits the same relationship, in the relationship of Figure 2b, the occurrence of points that exhibit a low value for *y* regardless of the value of *x* means that a true tradeoff does not exist. (The same argument can be extended to relationships like that in Figure 2c.) Furthermore, while a significant linear relationship can be fit to such triangular relationships (solid lines in Figure 2), they obscure the lack of relationship among the extreme quantiles of points (dashed and dotted lines in Figure 2). Only in hard tradeoffs (Figure 2a) are scaling slopes constant for all quantiles of the data. Harris and Pannell (2010) argue that there is a tradeoff between ramification and hydraulic efficiency, yet the bivariate distributions of these points is important to consider, as their interpretation hinges on whether the relationship more resembles that of Figure 2a or Figures 2b,c. If the latter, then the occurrence of individuals that can be both highly ramified and highly efficient would strongly question whether a tradeoff exists at all.

**Figure 2.**
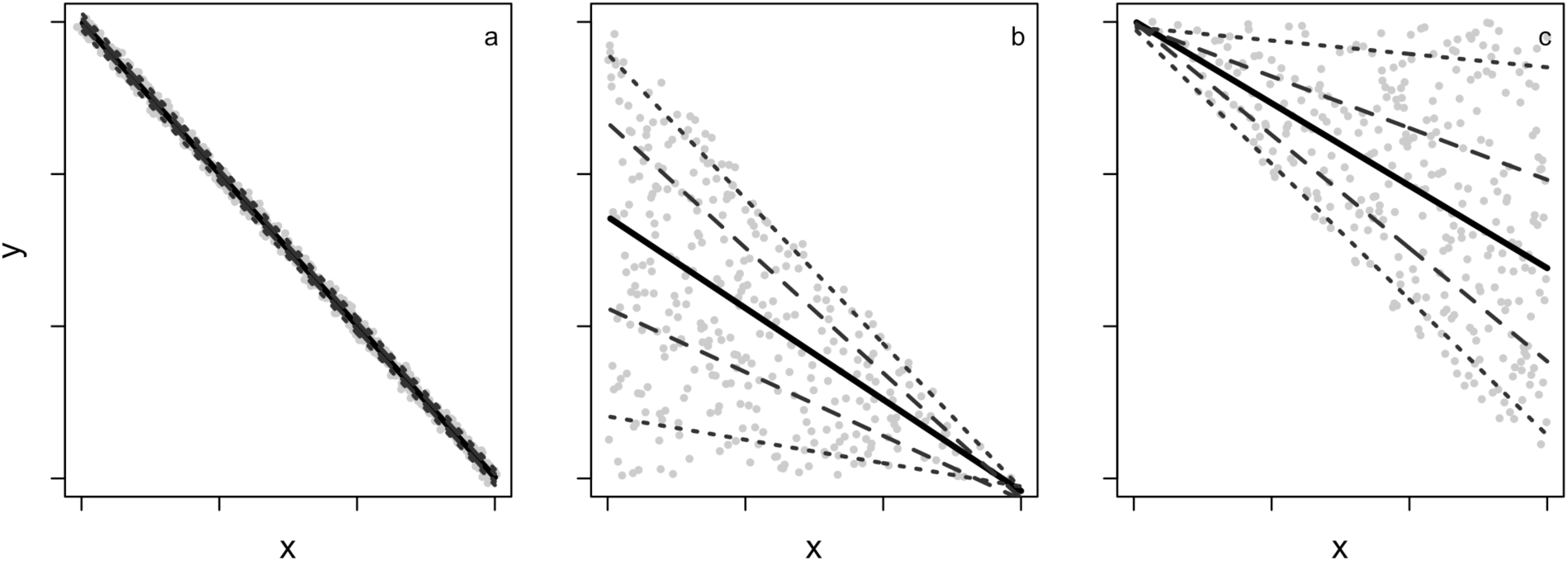
Three bivariate distributions of data considered to represent tradeoffs. In all panels, the solid line represents the regression through the median quantile, with dashed lines representing regressions through the 30% and 70% quantiles, and dotted lines representing regressions through the 10% and 90% quantiles. (a) A hard tradeoff in which all quantiles exhibit the same relationship (similar slopes but differing intercepts). (b,c) Although the bivariate distribution of points is bounded by an upper or lower limit and the median quantile exhibits a tradeoff, this tradeoff does not exist among all quantiles. Such relationships offer little support for the existence of a tradeoff.

In this study, we formally consider sexual differences in shoot hydraulic architecture and efficiency in two co-occurring *Leucadendron* species that differ dramatically in their degrees of sexual dimorphism. *L. daphnoides* is monomorphic with males and females that have similar degrees of ramification and leaf size, while *L. rubrum* is extremely dimorphic with males being more highly ramified with smaller leaves than females (Figure 1). We tested (1) whether hydraulic efficiency (per unit stem cross-sectional area or per unit shoot leaf area) scales with stem size (stem cross-sectional area) and branch ramification, (2) whether this relationship represents a hard tradeoff (i.e. Figure 2a), and (3) whether this differed between males and females within a species. We characterized branch ramification using the method of Harris and Pannell (2010), as well as a simpler method we developed for this study. We first compared these two metrics of ramification in the context of Corner’s rules (Corner 1949), with the expectation that to conform to Corner’s rules, a metric of ramification should scale predictably with leaf size. Our analyses suggest that hydraulic efficiency did not differ significantly betwene males and females within a species nor between the two co-occurring and intermixed species we examined. We conclude that sexual dimorphism in plant architecture requires alternate explanations than differences in hydraulic efficiency.

## Methods

### Plant material

Males and females of both species grow naturally near Du Toitskloof Pass, Western Cape, South Africa. All individuals were co-occurring within ~10m of each other on a slope extending westward toward the city of Paarl. The two species differ in the degree of serotiny, with *L. rubrum* holding its cones for an average of 2.8 years and *L. daphnoides* an average of 0.9 years (Williams 1972; Harris & Pannell 2010). Plants were collected and measurements made during November-December 2012 and during April-May 2013. In the field, shoots were cut at the plant base with garden shears and immediately recut under water at least one node above the previous cut. Individual shoots were placed in dark, plastic bags, and their bases submerged in water during transport back to the lab. Shoots were stored in a 4°C refrigerator until the day of measurement, and any shoots not measured within three days were discarded.

### Measurement of hydraulic conductance

Immediately prior to hydraulic measurements, shoots were defoliated underwater. When necessary, a sharp blade was used to cut leaves at the petiole base. Stems were recut underwater with a new, carbon steel blade. Hydraulic conductance was measured on whole, branching networks of various sizes using a low pressure flow meter (Kolb, Sperry & Lamont 1996). With this method, the cut stem base was inserted into a compression fitting (Omnifit) that was connected to a hard-sided tubing, the opposite end of which was submerged in a vial of filtered 0.01 M KCl that sat on a balance (Mettler-Toledo MS205DU with a resolution of 0.01 mg). The branched shoot was then placed inside a chamber connected to a vacuum pump. Flow rates from the balance were recorded automatically every 5-20 seconds (the frequency depended on the absolute flow rate) at each of a series of pressures below ambient: 10, 30, 50, 65, 40, 20 kPa. The stable flow rate at each pressure was determined when the coefficient of variation of the previous ten instantaneous flow rates was less than 5%. Hydraulic conductance, K, was obtained by a linear regression of flow rate as a function of pressure (kg s^−1^ MPa^−1^). *K* was normalized by either entire shoot leaf area (*K*_*LA*_) or by cross-sectional area of the stem base (*K*_*CSA*_).

### Quantifying ramification and other traits

At the end of each measurement, shoots were assessed for other morphological traits. The cross-sectional area of the stem base was calculated from the average of two perpendicular measurements of stem diameter at the base using manual calipers. Shoot ramification was quantified in two ways. First, we quantified the rate of diameter change down the length of a subset of the shoots, using the method of Harris and Pannell (2010; termed here ‘HP ramification’) and described briefly here. Starting at the highest branch, we measured stem cross-sectional area at the midpoint of each consecutive internode down the shoot as well as the relative position of this midpoint along the length of the shoot. The slope of the relationship between the logarithm of stem cross-sectional area and the relative distance from the crown provides an index of ramification. This method of quantifying ramification assumes that all branches arise immediately below terminal inflorescences and at nodes. For some species of *Leucadendron,* including *L. daphnoides,* this assumption is valid. However, in other species, particularly males of *L. rubrum,* most branches occur along the internodes. To account for these many small branches we measured the total number of branch tips on a shoot (regardless of their size and position on the shoot) normalized by the cross-sectional area of the stem base. The number of branch tips per stem cross-sectional area (BTSA) can easily be applied to shoots of any size, including shoot segments from the most recent year of growth that do not have properly defined internodes.

Leaves were placed on a flatbed scanner and their total area and number determined per shoot using ImageJ (Rasband 2012). The average leaf size per shoot (termed here simply ‘leaf size’) was determined by dividing the total leaf area by the total number of leaves on the shoot. For consistency with Corner’s rules, we quantified the average leaf area per branch by dividing the total shoot leaf area by the total number of terminal branches on each shoot.

Sexual dimorphism in each trait was calculated according to Harris and Pannell (2010):

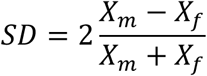

where X_m_ and X_f_ are mean trait values for males and females, respectively. This metric of quantifying the degree of sexual dimorphism is controversial, and we report a more accurate metric of dimorphism as (Lovich & Gibbons 1992):

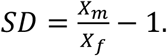

### Data analysis

All data were analyzed using R v. 3.0.2 (Team 2012). Scaling relationships were determined using standard major axis regression (SMA) as implemented in the ‘smatr’ package. For each pair of variables, relationships were determined for each species and sex combination. For certain scaling relationships, the slope test included in the function *sma* was used to test whether slopes significantly differed from unity. In some cases, data were pooled within species or across all species and sex combinations if there were no significant differences between the slopes of each group. Quantile regression was used to determine whether the relationship among the median quantile was consistent among all quantiles of the data, as implemented in the ‘quantreg’ package.

## Results

### Stem and leaf architectural relationships

The two study species differed dramatically in their degrees of sexual dimorphism in both leaf and stem traits (Figure 1, Table 1). The two metrics of branch ramification did not covary linearly, even in log-log space, and HP ramification showed less variability than did BTSA, with most individuals clumped around unity (Figure 3). HP Ramification estimated larger differences between males and females of the monomorphic species *L. daphnoides* than it did between males and females of the dimorphic species *L. rubrum.* It erroneously estimated *L. rubrum* females to be more highly ramified than males (Table 1).

**Figure 3.**
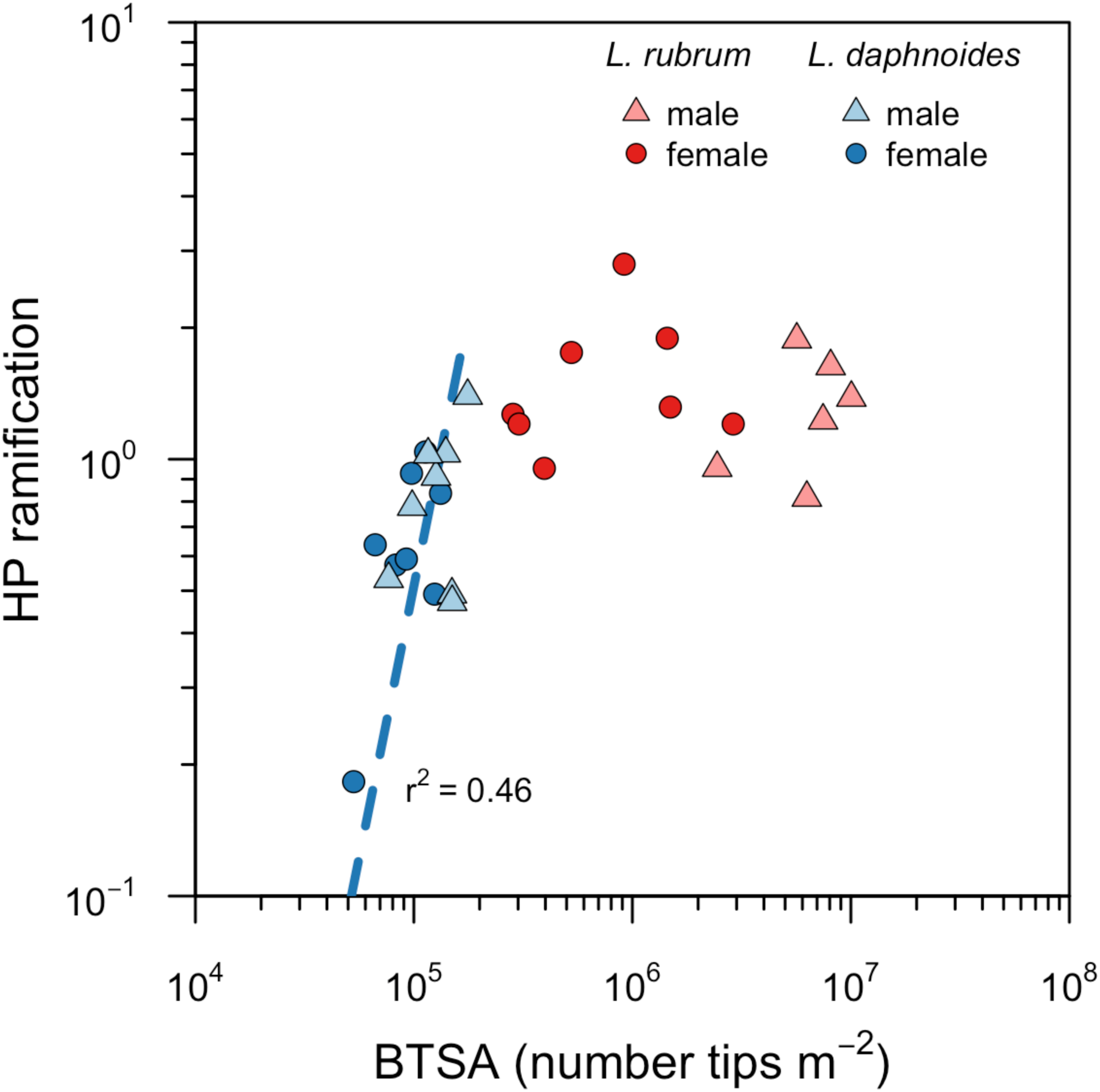
Relationship between the two metrics of ramification: that used by Harris and Pannell (2010; ‘HP ramification’) and that used in the present study (BTSA). Dashed line indicates significant relationship within species.

**Table 1.** Sexual dimorphism in morphological traits for the two study species, using two ways of quantifying dimorphism. Consistent with Harris and Pannell (2010), positive values of dimorphism indicate that males have higher trait values

| Trait | *Harris and Pannell*<br>*(2010)* |  | *Lovich and Gibbons*<br>*(1992)* |  |
| --- | --- | --- | --- | --- |
|  | *L. daphnoides* | *L.*<br>*rubrum* | *L. daphnoides* | *L. rubrum* |
| HP ramification | 0.65 | -0.16 | 0.75 | -0.15 |
| BTSA | 0.40 | 1.57 | 0.41 | 8.29 |
| Leaf size | -0.11 | -0.89 | 0.00 | -0.63 |
| Leaf area per tip | -0.39 | -1.68 | -0.29 | -0.92 |
| Vessel area | 0.06 | -1.19 | 0.03 | -0.74 |
| Leaf area:sapwood<br>area | 0.03 | 0.27 | 0.07 | 0.40 |

Further support for BTSA as a better metric of ramification was found in the scaling relationships based on Corner’s rules (Figure 4). Corner’s rules predict that leaf size, branch size, and branch ramification should be allometrically related traits. Leaf area per branch tip scaled isometrically with BTSA (R^2^ = 0.93, *P* < 0.001; slope = -0.97, *P*_α=1_ = 0.41; Figure 3b) but not with HP ramification, for which a weak, significant relationship existed only for *L. daphnoides* (*R*^2^ = 0.43, *P* = 0.005; Figure 3a). Corner’s rules predict that leaf size should scale with branch ramification, yet there was no relationship between leaf size and HP ramification. However there were highly significant, species-specific scaling relationships between leaf size and BTSA *(L. daphnoides: R*^2^ = 0.46, *P* < 0.001; *L. rubrum: R*^2^ = 0.87, *P* < 0.001). Only relationships predicted by Corner’s rules were statistically significant at all quantiles both within and among species (Table 2), suggesting that these relationships are likely hard tradeoffs.

**Figure 4.**
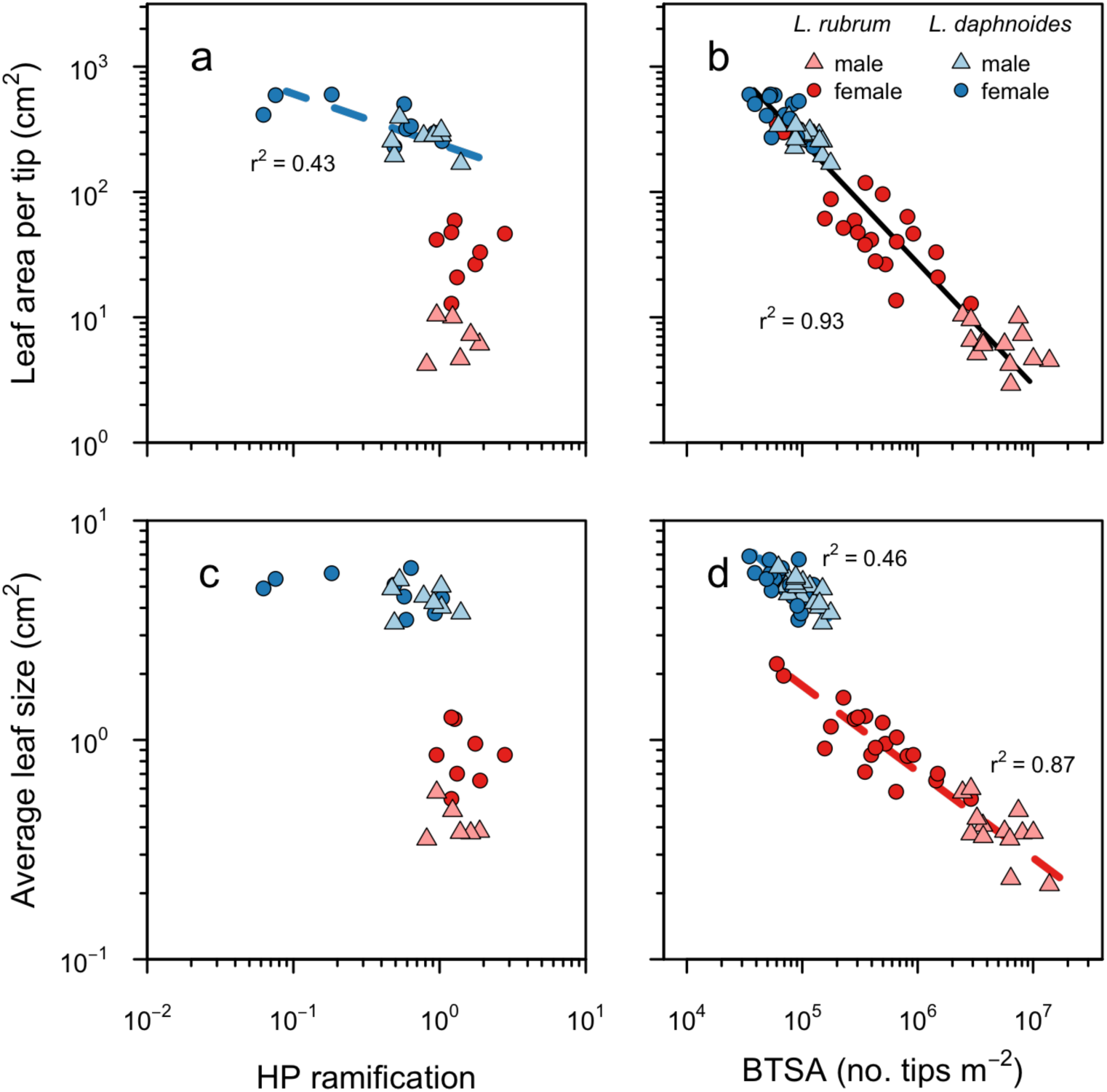
Corner’s rules and the relationships between branch ramification (BTSA), leaf size, and leaf area per branch tip. Solid lines indicate statistically significant relationships across all species x sex combinations (P < 0.001). Dashed lines indicate significant relationships within species (P < 0.001). (a) Leaf area per tip as a function of HP ramification. (b) Relationship between leaf area per tip and BTSA. This scaling slope was highly significant (R^2^ = 0.93, P < 0.001) across species and sexes with a slope of -0.974 (-1.039, -0.913) and was not significantly different from isometry (P = 0.41). (c) Relationship between leaf size and HP ramification. (d) Relationship between leaf size and BTSA. These scaling slopes were highly significant within species *(L. daphnoides:* slope = -0.453 (-0.588, -0.349), R^2^ = 0.46, P < 0.001; *L. rubrum:* slope = -0.392 (-0.448, -0.344), R^2^ = 0.87, P < 0.001) and were not significantly different from each other (P = 0.33).

### Hydraulic efficiency and shoot size

Leaf-area normalized shoot hydraulic conductance, *K*_*LA*_, decreased with whole shoot leaf area for *L. rubrum* females (*R*^2^ = 0.29, *P* = 0.02) and for *L. daphnoides* males (*R*^2^ = 0.59, *P* < 0.001; Figure 5a), but there was no relationship between shoot leaf area and *K*_*LA*_ for *L. daphnoides* females and *L. rubrum* males. *K*_*LA*_ also decreased with stem cross-sectional area for *L. daphnoides* males (*R*^2^ = 0.59, *P* < 0.001), but not for the other groups (Figure 5b). Hydraulic conductance normalized by stem cross-sectional area, *K*_*CSA*_, decreased with increasing shoot leaf area for *L. daphnoides* males (*R*^2^ = 0.34, *P* = 0.01), but not for the other groups (Figure 5c). Contrary to the predictions of Harris and Pannell (2010), for both *L. rubrum* females (*R*^2^ = 0.28, *P* < 0.001) and *L. daphnoides* males (*R*^2^ = 0.52, *P* = 0.001), *K*_*CSA*_ decreased with stem cross-sectional area, but not for *L. rubrum* males or *L. daphnoides* females (Figure 5d).

**Figure 5.**
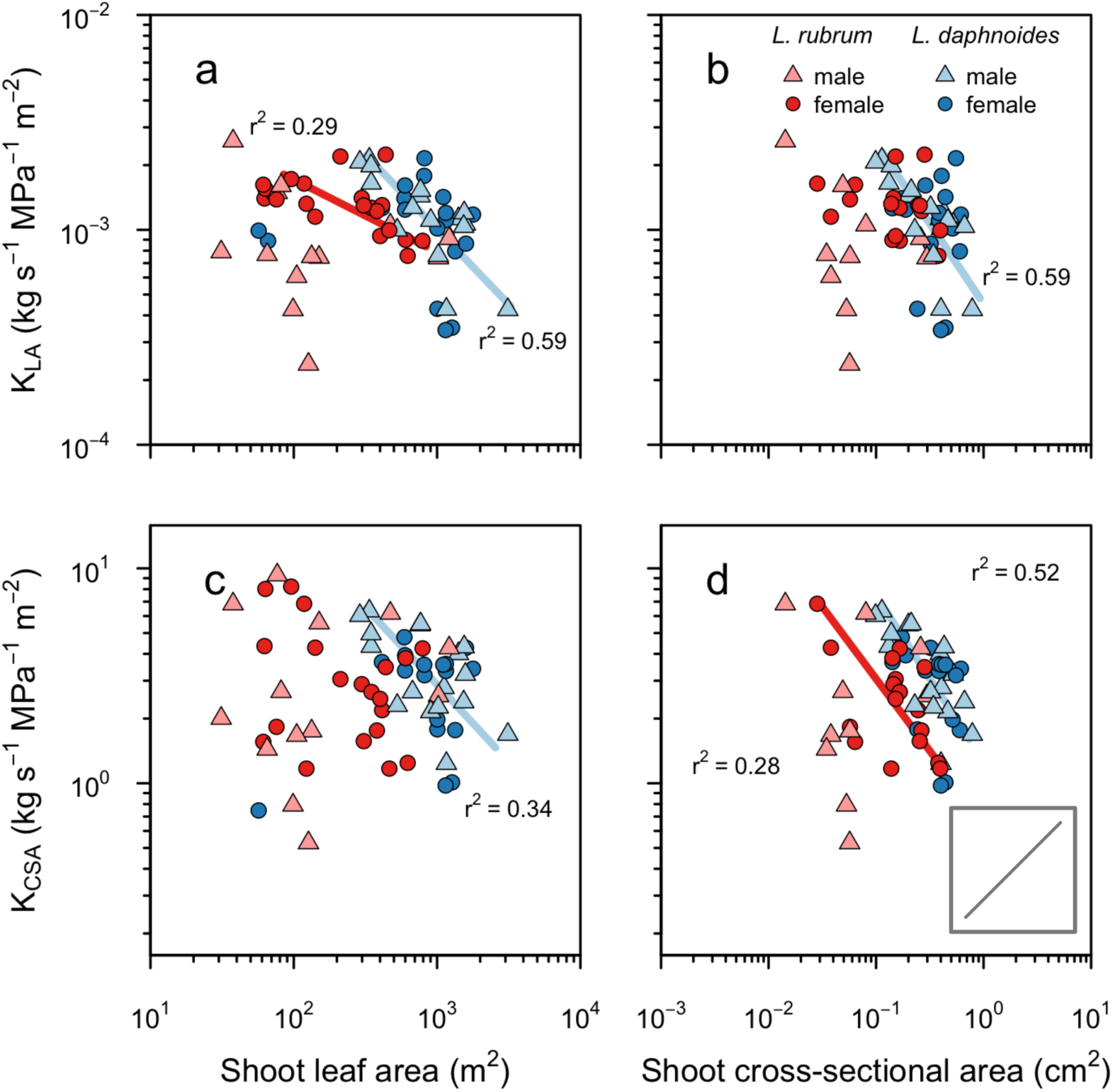
The relationships between two measures of shoot hydraulic efficiency and two measures of shoot size. Lines indicate statistically significant relationships of sex x species combinations (all P < 0.05). Inset indicates predictions made by Harris and Pannell (2010). Relationships between shoot hydraulic conductance normalized by shoot leaf area and (a) shoot leaf area and (b) stem cross-sectional area. Relationships between shoot hydraulic conductance normalized by stem cross-sectional area and (c) shoot leaf area and (d) stem cross-sectional area.

### Hydraulic conductance and branch ramification

Neither way of quantifying ramification had any consistent effect on *K*_*CSA*_ (Figure 6a,b). The only effect of ramification on *K*_*CSA*_ occurred in *L. daphnoides* females, but this negative relationship was driven solely by the inclusion of three current year shoots lacking any nodes and having but a single branch tip (circles points in Figure 6b). For *L. daphnoides* females *K*_*LA*_ decreased with increasing BTSA *(R*^2^ = 0.36, *P* = 0.006), and pooling *L. rubrum* males and females together showed a similar negative relationship between *K*_*LA*_ and BTSA for this species (*R*^2^ = 0.16, *P* < 0.001), but not for *L. daphnoides* males (Figure 6c). Quantile regression revealed that lower quantiles exhibited stronger, more statistically significant slopes than higher quantiles for both species (Table 2), consistent with the relationship exhibited in Figure 2c. There was no relationship between *K*_*LA*_ and HP ramification (Figure 6d).

**Figure 6.**
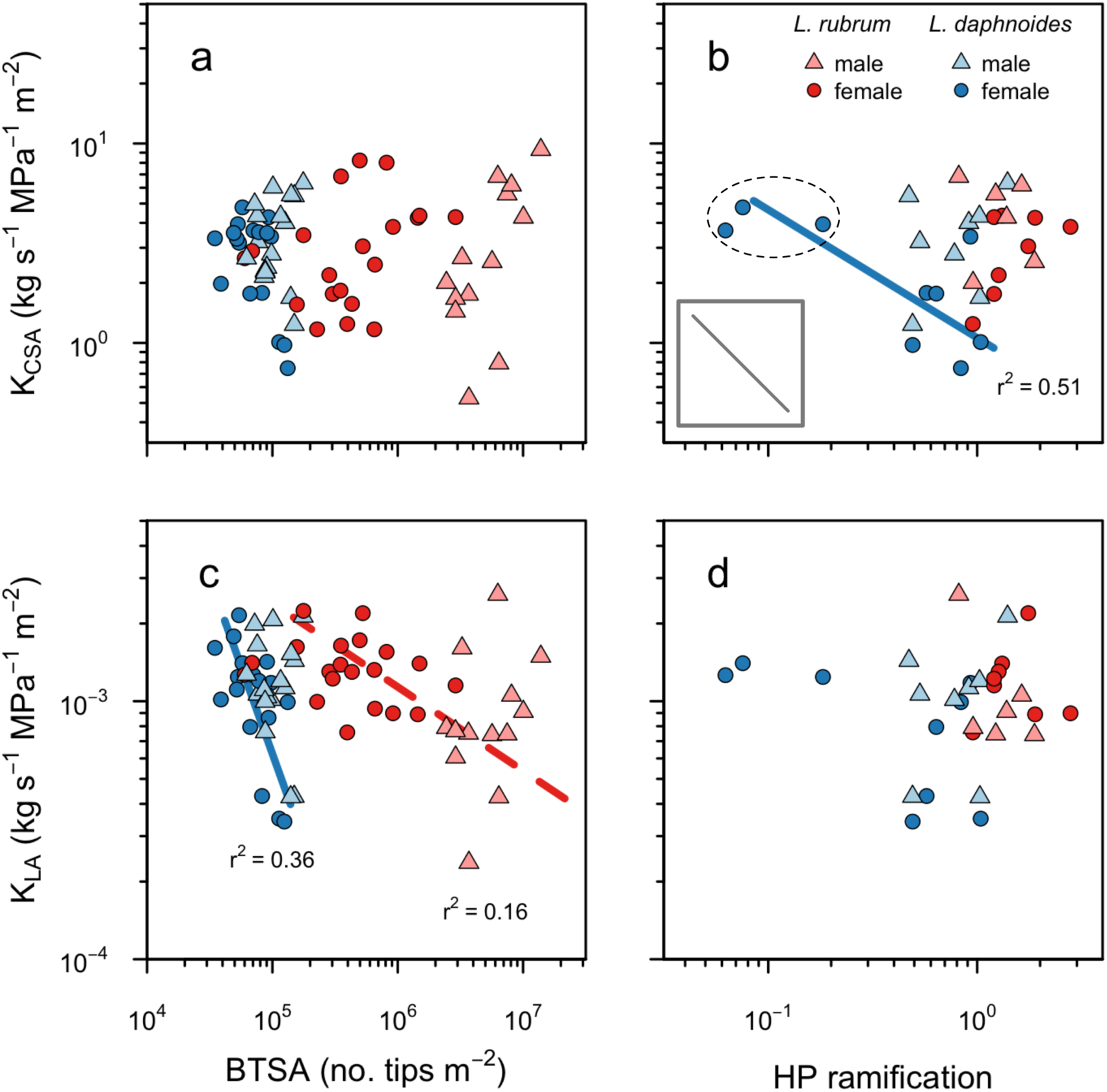
The relationships between two measures of hydraulic conductance and two measures of branch ramification. (a) Shoot hydraulic conductance per unit cross-sectional stem area as a function of BTSA. (b) Shoot hydraulic conductance per unit cross-sectional stem area as a function of HP ramification. The three circled points driving the relationship for *L. daphnoides* females are shoots from a single year’s growth that lacked an internode. Without these points, there is no significant relationship. (c) Shoot hydraulic conductance per unit leaf area as a function of BTSA. (d) Shoot hydraulic conductance per unit leaf area as a function of HP ramification. In (c), solid line indicates a significant relationship within a sex x species combination, and dashed line indicates a significant relationship within a species (P < 0.01). Inset indicates predictions made by Harris and Pannell (2010).

## Discussion

In the present analysis of two co-occurring *Leucadendron* species differing in their degrees of sexual dimorphism, we found no consistent support for hydraulic differences between the sexes. Although measurements on only two species preclude determining the effects of serotiny on shoot hydraulics, many of the predictions proposed by Harris and Pannell (2010) should exist both within and among species. Yet we found little support for their predictions–and, indeed, in one case a pattern opposite to their prediction–particularly for the metric of ramification used in their analysis and as the foundation for their arguments. Our results, therefore, question the relationship between sexual dimorphism in branch ramification and serotiny reported by Harris and Pannell (2010). Their results depend, first, on accurately quantifying branch ramification and, second, on hydraulic efficiency varying as a function of shoot size and ramification.

Accurately characterizing ramification in ways that reflect obvious differences between sexes and species and that conform to known scaling relationships is fundamental to determining how ramification, and sexual dimorphism in it, may be related to other traits. The method used by Harris and Pannell (2010) for quantifying shoot ramification did not accurately describe obvious, qualitative differences between sexes and species (Figure 1), was not related to another metric of ramification (BTSA; Figure 3), and, most importantly, did not characterize known scaling relationships between ramification and leaf size that have been supported by decades of theory and data (Figure 4). Corner’s rules predict that “the stouter the stem, the bigger the leaves and the more complicated their form,” and, more apropos to the present study, “the greater the ramification, the smaller become the branches and their appendages” (Corner 1949). Corner’s first prediction between stem size and leaf size has been well-characterized for many species including *Leucadendron* (Bond & Midgley 1988; Ackerly & Donoghue 1998; Olson *et al.* 2009). Corner’s second predictions, however, has been less well-studied. Using the metric of ramification proposed by Harris and Pannell (2010), we found no consistent relationship across species and sexes between HP ramification and either leaf area per branch tip or leaf size (Figure 4a,c). However, leaf area per terminal branch tip did scale isometrically with BTSA with a single slope describing the relationship across species and sexes, as predicted by Corner’s rules, and allometrically with leaf size with species-specific slopes and intercepts (Figure 4b,d). The metric of ramification used by Harris and Pannell (2010), therefore, fails to accurately quantify branch ramification in two important ways. It predicted greater dimorphism in ramification for the monomorphic *L. daphnoides* than for the obviously dimorphic *L. rubrum* (Figures 2,3; Table 1), and it does not conform to well-established scaling relationships (Figure 4). Furthermore, only the relationships predicted by Corner’s rules were significant at all quantiles, suggesting they reflect strict, biophysical tradeoffs that define fundamental axes of plant architecture (Table 2, Figure 2a).

Shoot hydraulic conductance did increase with shoot size, whether quantified by whole shoot leaf area or stem cross-sectional area (data not shown), but this relationship is not at all surprising because larger shoots should move absolutely more water. However, once hydraulic conductance was normalized by shoot size–either total shoot leaf area or stem cross-sectional area–there was no increase in size-normalized hydraulic conductance with shoot size (i.e. no increase in hydraulic efficiency with shoot size; Figure 5). In fact, the opposite pattern was found. Both *K*_*LA*_ and *K*_*CSA*_ decrease with increasing shoot size (Figure 5). Contrary to the predictions of Harris and Pannell (2010), large stems are no more hydraulically efficient than small stems. Thus, larger stems, whether males or females, provide no more water to their transpiring leaves per unit leaf area or stem cross-sectional area than do smaller stems. Indeed, the opposite is true. This hydraulic limitation may determine maximum shoot size and whether additional growth at the plant level is apportioned to growing an individual shoot or to adding new shoots, as it does for root growth (Bouda, Brodersen & Saiers in review).

Second, Harris and Pannell (2010) assumed that more highly ramified shoots would be less hydraulically efficient. In other words, they predicted that size-normalized shoot hydraulic conductance should decrease with increasing ramification. On this point, we found mixed results. Harris and Pannell (2010) focused exclusively on hydraulic conductance per unit stem cross-sectional area (*K*_*CSA*_) as their preferred metric of hydraulic efficiency. We found that *K*_*CSA*_ was largely unrelated to branch ramification, either assessed using their metric of ramification or BTSA (Figure 6a,b). Only for *L. daphnoides* females was *K*_*CSA*_ negatively correlated with HP ramification, but this relationship was driven by three small shoots that did not include any nodes (circled points in Figure 6b). Excluding these three shoots eliminated any significant relationship between HP ramification and *K*_*CSA*_ either within or among groups. However, we did find a significant negative relationship (median quantile regression) between leaf area-normalized shoot hydraulic conductance (*K*_*LA*_) and ramification for *L. daphnoides* females and for *L. rubrum* once males and females were pooled together (Figure 6c). However, quantile regression provided no consistent support for this relationship (Table 2). In fact, the range of *K*_*LA*_ for the more highly ramified *L. rubrum* males was greater than that of the conspecific females (Figures 6a,c), suggesting that highly ramified males are capable of transporting water as efficiently or more efficiently than less ramified females. Thus, compelling evidence for a hard tradeoff between shoot ramification and hydraulic efficiency is lacking in our data, the distribution of which more resembles the relationship shown in Figure 2c than that in Figure 2a. Furthermore, there is more variation in sexual dimorphism among weakly serotinous *Leucadendron* species (those that hold their cones for one year or less) than there is among all other species that maintain their cones for longer than one year, further suggesting that there is no hard tradeoff (sensu Grubb 2016) between serotiny and branch ramification (Figure 3 in Harris and Pannell 2010).

We found no compelling evidence that hydraulic efficiency differs between conspecific males and females or even between the co-occurring species we studied here, which differ dramatically in their degrees of sexual dimorphism. Males and females of each species must survive to the next fire events in order to be represented in the most recent year’s seed crop and, thus, maximize fitness (Midgley 2000). Conspecific males and females should, therefore, have similar longevities and likely also similar rates of metabolism. This is supported by the similarity in hydraulic efficiencies among conspecifics we studied and the strong correspondence between shoot hydraulic efficiency and photosynthetic capacity (Brodribb & Feild 2000; Brodribb, Holbrook & Gutiérrez 2002). If the hydraulic architecture of females were more efficient than that of males, then it is not clear why males would not also have evolved a similar branching and leaf structure given that males and females co-occur and may compete with each other. Also, if the hydraulic architecture of males were less efficient, then males may incur costs associated with reproduction equal to or higher than those of females, contrary to most theory and data regarding the costs of reproduction (Bond & Maze 1999). Instead, our data and those of Midgley (2010) are consistent with there being similar hydraulic efficiencies between species and sexes, regardless of the degrees of sexual dimorphism and branch ramification.

What, then, may be driving the pattern between sexual dimorphism in branch ramification and the degree of serotiny observed by Harris and Pannell (2010)? Because sexual dimorphism is a difference between males and females, it may arise from selection on just males, on just females, or on divergent selection on both males and females. Thus, sexual dimorphism may be due to multiple factors that all contribute to sexual dimorphism to varying degrees. The present study suggests that differences in shoot ramification cannot be explained by differences in the hydraulic costs of reproduction. The occurrence of many highly dimorphic, non-serotinous species in the genus (e.g. *L. ericifolium* and *L. pubescens;* Figure 3 in Harris and Pannell 2010) suggests that dimorphism in shoot ramification and related traits is unlikely to be explained solely by serotiny. One untested mechanism, though, is based on differences in the biomechanical requirements of male and female shoots. Females of all *Leucadendron* species must bear cones, at least for several months even in non-serotinous species, and this biomechanical constraint may mean that females have thicker stems, which may be exacerbated by increased serotiny. Because flexural stiffness increases with the fourth power of stem radius (Vogel 2013), selection for thicker stems would have a dramatic effect on stem stiffness. Producing thicker stems may not be very costly if increases in stem diameter are driven predominantly by additional pith in the middle of the stem or even as new sap wood is laid down. Corner’s rules have been shown to extend to the cone size-branch size relationship among conifers (Leslie *et al.* 2014), and there may be a similar relationship within *Leucadendron,* as there is with inflorescences (Midgley & Bond 1989). At the same time, males are under selection for increased ramification to support more inflorescences that must be highly flexible, particularly in wind-pollinated species (Midgley & Bond 1989; Welsford, Midgley & Johnson 2014). These differing biomechanical constraints between the sexes may underlie sexual dimorphism and allow for an emergent relationship between shoot ramification and serotiny. Yet, the magnitude of sexual dimorphism may be mitigated by differing degrees of developmental constraints within species, which would weaken the effects of selection on shoot biomechanics and, potentially, lead to a weak relationship between serotiny and dimorphism in ramification like that characterized by Harris and Pannell (2010). It is unlikely that a single predictor may explain something as complex as sexual dimorphism.

